# Differential homeostatic regulation of glycinergic and GABAergic nanocolumns at mixed inhibitory synapses

**DOI:** 10.1101/2020.11.23.372383

**Authors:** Xiaojuan Yang, Hervé Le Corronc, Pascal Legendre, Antoine Triller, Christian G Specht

## Abstract

Super-resolution imaging of synapses has revealed that key synaptic proteins are dynamically organized within sub-synaptic domains (SSDs). At mixed inhibitory synapses in spinal cord neurons, both GlyRs and GABA_A_Rs reside at the same post-synaptic density (PSD). To examine how the different inhibitory receptors are organized and regulated, we carried out dual-color direct stochastic optical reconstruction microscopy (dSTORM). We found that endogenous GlyRs and GABA_A_Rs as well as their common scaffold protein gephyrin form SSDs that align with pre-synaptic RIM1/2, thus forming trans-synaptic nanocolumns. Strikingly, GlyRs and GABA_A_Rs occupy different sub-synaptic spaces, exhibiting only a partial overlap at mixed inhibitory synapses. When network activity was increased by pharmacological treatment using the K^+^ channel blocker 4-aminopyridine (4-AP), the GABA_A_R copy numbers of as well as the number of GABA_A_R SSDs were reduced, while GlyRs remained largely unchanged. This differential regulation is likely the result of changes in gephyrin phosphorylation that preferentially occurred outside of the SSDs. The total gephyrin content was not altered by 4-AP application. The activity-dependent regulation of GABA_A_Rs versus GlyRs suggests that different signaling pathways control their respective sub-synaptic organization. Whereas gephyrin serves as a scaffold protein that upholds GlyR numbers at SSDs, it may act as a switch regulating GABA_A_Rs via its phosphorylation state. Taken together, our data reinforce the notion that the precise sub-synaptic organization of GlyRs, GABA_A_Rs and gephyrin has functional consequences for the homeostatic regulation of mixed inhibitory synapses.

**Highlights:** Alignment of sub-synaptic domains (SSDs) in trans-synaptic nanocolumns at inhibitory synapses Differential spatial organization of SSDs formed by GlyRs and GABA_A_Rs at mixed inhibitory synapses Activity-dependent regulation of GABA_A_Rs but not GlyRs at mixed inhibitory synapses Gephyrin phosphorylation is compartmentalized in SSDs within the synaptic scaffold

## Introduction

The application of single molecule localization microscopy (SMLM) has provided new information about the nanoscale organization of synapses. Numerous post-synaptic proteins are heterogeneously distributed at synapses and compartmentalized into sub-synaptic domains (SSDs) (discussed in (Yang & Specht, 2019)). At excitatory synapses, AMPARs assemble dynamically in SSDs that are stabilized by the scaffold protein PSD-95 (MacGillavry et al., 2013, Nair et al., 2013). At inhibitory synapses, gephyrin molecules also form SSDs, which are re-organized during inhibitory synaptic plasticity (Pennacchietti et al., 2017, Specht et al., 2013). Furthermore, Tang and colleagues have shown that post-synaptic SSDs at excitatory synapses are aligned with pre-synaptic SSDs, forming so-called trans-synaptic nanocolumns (Tang et al., 2016). This led to the proposal that sub-synaptic domains are functional units underlying synaptic plasticity.

Glycine receptors (GlyRs) and gamma-aminobutyric acid type A receptors (GABA_A_Rs) are the two main inhibitory receptors mediating fast inhibition in the central nervous system. In the spinal cord, GlyRs and GABA_A_Rs co-exist at the same post-synaptic density (PSD) during development and into adulthood (Chery & de Koninck, 1999, Dumoulin et al., 2000, Legendre, 2001, Todd et al., 1996, Triller et al., 1987). These synapses are referred to as mixed inhibitory synapses. The corresponding neurotransmitters, glycine and GABA, are co-released from the same pre-synaptic vesicles (Jonas et al., 1998, Keller et al., 2001, O’Brien & Berger, 2001). Glycinergic and GABAergic mIPSCs have distinct kinetics, the latter exhibiting slower decay times (Alvarez, 2017). Moreover, previous studies have shown that the lateral diffusion of synaptic GlyRs and GABA_A_Rs are differentially regulated at mixed inhibitory synapses by excitatory activity and microglial-dependent signaling (Cantaut-Belarif et al., 2017, Levi et al., 2008). These observations raise the possibility that the selective control of GlyRs and GABA_A_Rs could underlie the plasticity of mixed inhibitory synapses.

Gephyrin is the main scaffold protein at inhibitory synapses and a key player for both GlyR and GABA_A_R clustering. Biochemical studies have shown that gephyrin molecules form stable trimers through interactions of their G-domains, and that they can dimerize via their E-domains (Saiyed et al., 2007, Schrader et al., 2004, Sola et al., 2004). It has thus been proposed that gephyrin molecules form a hexagonal planar lattice below the post-synaptic membrane (Kneussel & Betz, 2000). Gephyrin provides binding sites for both GlyRs and GABA_A_Rs, even though the binding affinity for GlyRs is substantially higher than that of GABA_A_Rs (reviewed in (Tretter et al., 2012). The binding sites for the different receptor subunits in the gephyrin E-domain overlap, resulting in a competition between GlyRs and GABA_A_Rs for the same binding pocket of gephyrin at mixed inhibitory synapses {Marie, 2011 #1815). Furthermore, gephyrin molecules are prone to numerous post-translational modifications such as phosphorylation at multiple amino acid residues within the C-domain, which are important for the regulation of receptor binding and/or gephyrin clustering (Ghosh et al., 2016, Kalbouneh et al., 2014, Kuhse et al., 2012, Niwa et al., 2019, Tyagarajan et al., 2013, Tyagarajan et al., 2011).

In this study, we focused on the nanoscale organization of mixed inhibitory synapses in spinal cord neurons and investigated how endogenous GlyRs and GABARs are regulated at the sub-synaptic level using two-color dSTORM. Our data identify trans-synaptic nanocolumns comprising SSDs of RIM1/2, a component of the pre-synaptic active zone (AZ), as well as the post-synaptic scaffold protein gephyrin, GlyRs and GABA_A_Rs. Interestingly, the two types of receptor were partially segregated, each dominating different sub-synaptic domains. While GlyR clustering was insensitive to short-term changes in network activity, GABA_A_Rs were differentially regulated by neuronal activity, implying a key role in the homeostatic plasticity of mixed inhibitory synapses.

## Results

### Trans-synaptic nanocolumns at inhibitory synapses in spinal cord neurons

To probe the spatial relationship between pre- and post-synaptic elements at inhibitory synapses in spinal cord neurons, we conducted dual-color dSTORM imaging in native tissue. Semi-thin sections (2 μm) of mouse spinal cord tissue were immuno-labeled with antibodies against the post-synaptic scaffold protein gephyrin and the pre-synaptic AZ protein RIM1/2, and imaged by dSTORM (Figure 1). The gephyrin clusters in dSTORM images exhibited a heterogeneous distribution, forming sub-synaptic domains (SSDs) that are visible both in pointillist representations as well as rendered images (Figure 1A-C). Two-color dSTORM images of gephyrin and RIM1/2 further revealed a close association between gephyrin SSDs and RIM1/2 SSDs (Figure 1D). This alignment of pre- and post-synaptic SSDs is reminiscent of trans-synaptic nanocolumns that have been identified at excitatory synapses in primary cultured neurons (Tang et al., 2016). The identification of gephyrin SSDs and their alignment with pre-synaptic release sites *in vivo* points to a possible role for synaptic function.

**Figure 1.**
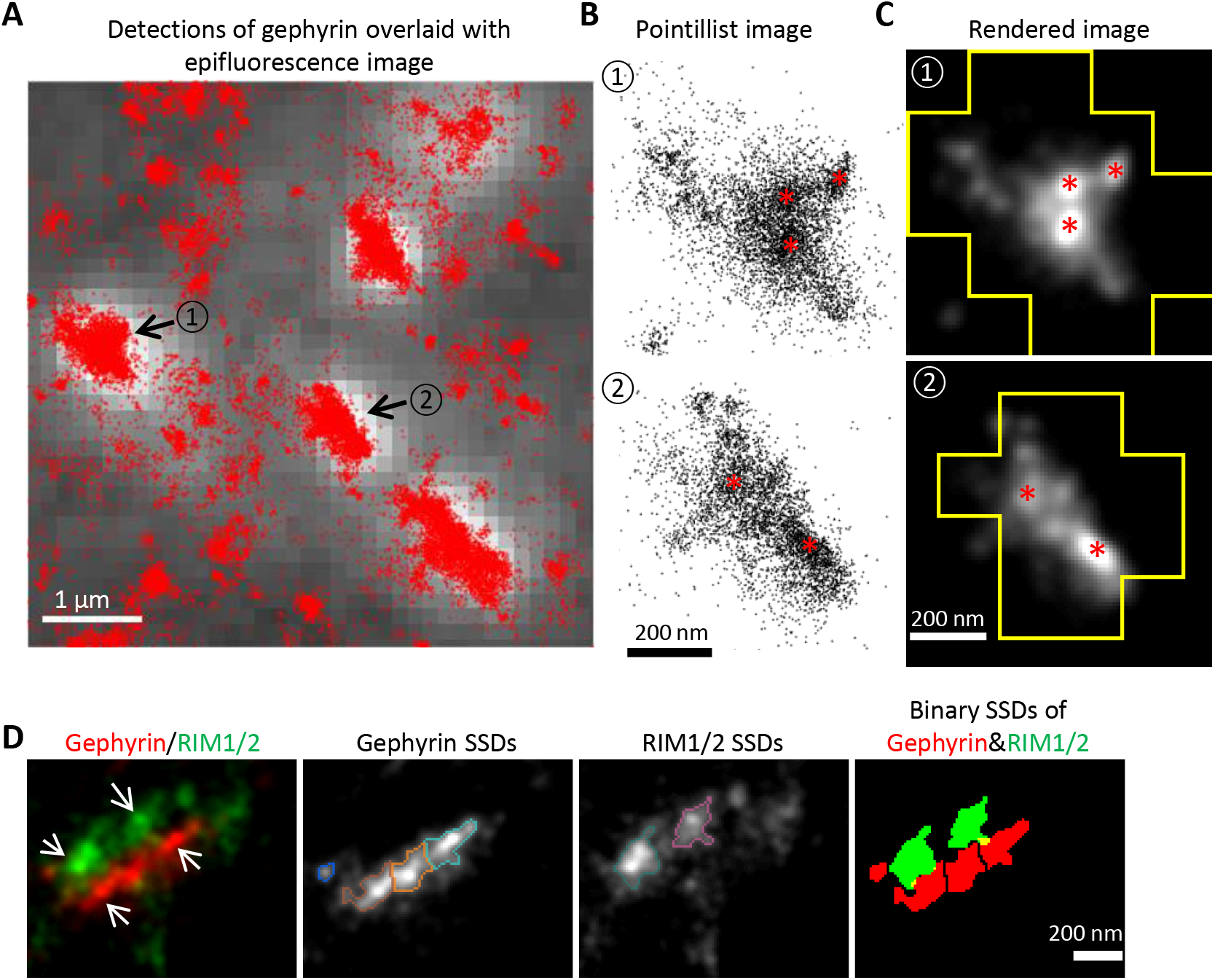
Gephyrin SSDs and their alignment with pre-synaptic RIM1/2 *in vivo*. (A-C) dSTORM imaging of gephyrin in sucrose impregnated cryosections of adult mouse spinal cord. (A) dSTORM detections of gephyrin (red dots) overlaid with the epifluorescence image (white with grey background). Synaptic clusters of gephyrin in dSTORM images were identified by the epifluorescence puncta. Scale bar: 1 μm. (B) Enlarged pointillist images of the two gephyrin clusters indicated in (A). (C) Rendered images of the two gephyrin clusters outlined with the boundaries of the epifluorescence mask. Gephyrin SSDs are indicated with red asterisks in (B) and (C). Scale bar: 200 nm. (D) Two-color dSTORM imaging of RIM1/2 and gephyrin in cryosections. From left to right: rendered images of RIM1/2 and gephyrin clusters showing the aligned SSDs (arrows); gephyrin SSDs segmented by watershed outlined with different colors; RIM1/2 SSDs outlined with different colors; binary SSDs of gephyrin and RIM1/2. Inhibitory synapses were identified by the gephyrin clusters in the epifluorescence images. Scale bar: 200 nm. RIM1/2 was labeled with Alexa 647, gephyrin (mAb7a) with Cy3B in these experiments.

For a closer characterization of the spatial organization we made use of cultured spinal cord neurons for our subsequent experiments. Two-color dSTORM imaging of synapses in cultured neurons was conducted on the target proteins in a pair-wise manner (see Methods). To quantify the pre- and post-synaptic organization along the trans-synaptic axis, super-resolution images of synapses with side view profiles were selected from the rendered dSTORM images. These images revealed a heterogeneous distribution of RIM1/2, gephyrin (antibody mAb7a), GlyRs and GABA_A_Rs, all of which formed distinctive SSDs at inhibitory synapses (Figure 2). The number of SSDs per synaptic cluster for these proteins was generally two to four, and the SSD counts increased with the synaptic cluster area (Figure S1). Importantly, post-synaptic SSDs of gephyrin, GlyRs and GABA_A_Rs were often aligned with pre-synaptic RIM1/2 SSDs (Figure 2).

**Figure 2.**
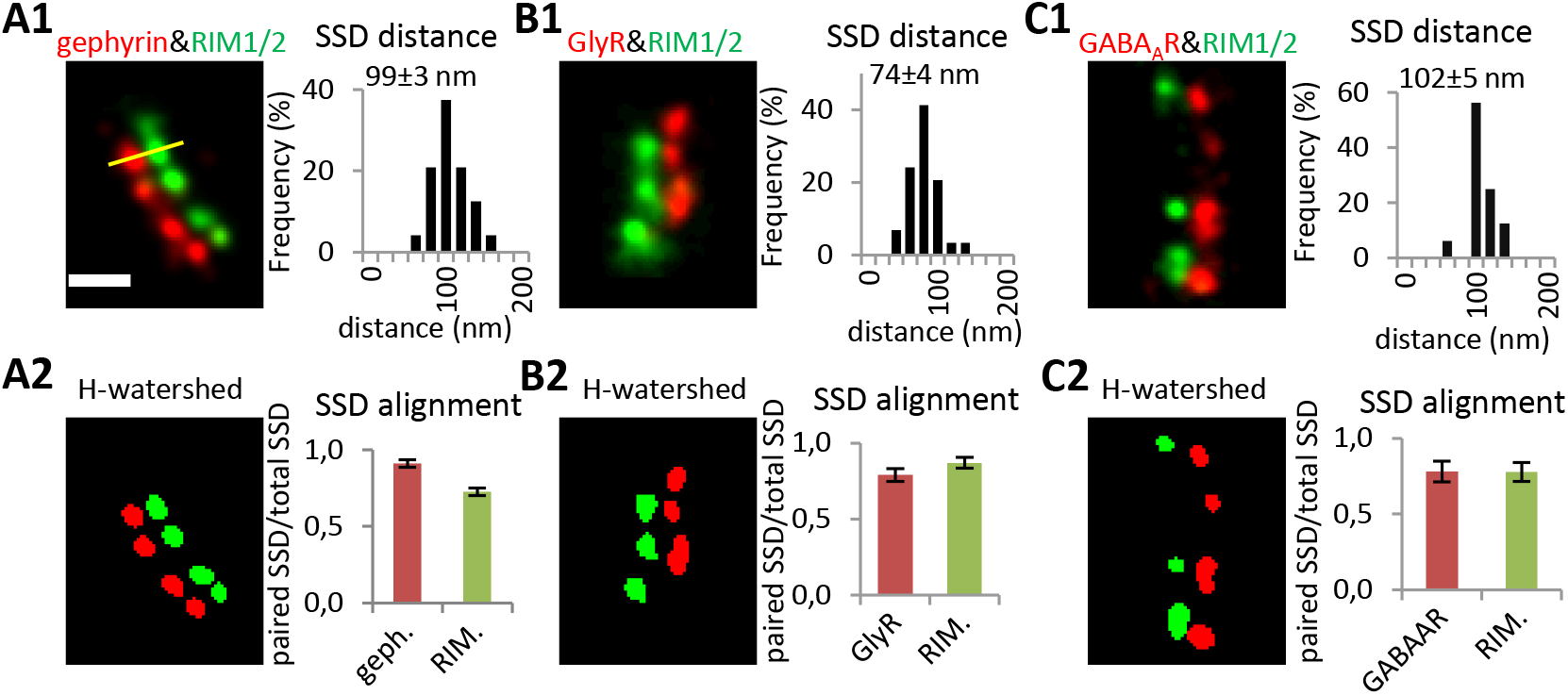
Trans-synaptic alignment at inhibitory synapses in cultured spinal cord neurons. (A1-C1) SSD distances between post-synaptic gephyrin, GlyRs, GABA_A_Rs and pre-synaptic RIM1/2, respectively. Distances were measured between the intensity peaks of paired SSDs in rendered dualcolor dSTORM images (yellow line, values given in mean ± SEM). RIM1/2 was labeled with Alexa 647 and gephyrin (mAb7a) with Cy3B in (A1). RIM1/2 was labeled with Cy3B, while GlyRs or GABA_A_Rs were labeled with Alexa 647 in (B1) and (C1). (A2-C2) Trans-synaptic alignment of SSDs of gephyrin, GlyRs, GABA_A_Rs and RIM1/2. SSDs were segmented by H-watershed. Number of synapses: n = 48 (A1, A2) from three independent experiments (no treatment), n = 30 (B1, B2) from two experiments with TTX treatment, n = 16 (C1, C2) from two experiments with TTX. Only synapses with side view profiles were included. Scale: 200 nm.

We measured the distances between the paired pre-synaptic SSDs and post-synaptic SSDs (Figure 2A1-C1, see Methods). The average distance between SSDs of pre-synaptic RIM1/2 and post-synaptic gephyrin and was 99 ± 3 nm (mean ± SEM, from 48 synapses). The distance between SSDs of RIM1/2 and GABA_A_Rs was very similar, measuring 102 ± 5 nm (from 16 synapses), however, the distance between RIM1/2 and GlyR SSDs was only 74 ± 4 nm (from 30 synapses). This difference is likely due to the fact that GlyRs were labeled using antibodies against an extracellular epitope, whereas GABA_A_Rs were labeled at an intracellular epitope. The apparent distance of 28 nm between the extracellular and the intracellular epitope is consistent with the longitudinal dimension of receptor complexes (~20 nm; (Patriarchi et al., 2018)) and the size of a single antibody (12-15 nm; (Maidorn et al., 2016)). These measurements also illustrate that the sequential immuno-labeling with classical antibodies does not add cumulative distances to the distance measurement, probably due to the random orientation of the antibodies.

Next, we quantified the degree of alignment between pre- and post-synaptic structures, expressed as the ratio of paired SSDs over the total SSDs for each target protein (Figure 2A2-C2). The majority of gephyrin SSDs were aligned with RIM1/2 SSDs (0.91 ± 0.03, mean ± SEM), and most RIM1/2 SSDs were in turn also aligned with SSDs of gephyrin (0.73 ± 0.04). Similarly, RIM1/2 SSDs were aligned with GlyR and GABA_A_R SSDs (0.87 ± 0.03 and 0.78 ± 0.06, respectively) and *vice versa* (0.79 ± 0.04 and 0.78 ± 0.07, respectively). These results confirm the existence of trans-synaptic nanocolumns at inhibitory synapses in cultured spinal cord neurons.

Trans-synaptic nanocolumns consisting of RIM1/2 and gephyrin SSDs may recruit both GlyRs and GABA_A_Rs at mixed inhibitory synapses. This is consistent with a recent study using STED and SIM super-resolution imaging, showing that gephyrin SSDs as well as GABA_A_R SSDs were closely associated with RIM1/2 SSDs at hippocampal and cortical synapses (Crosby et al., 2019). Given that trans-synaptic nanocolumns were also observed at excitatory synapses (Haas et al., 2018, MacGillavry et al., 2013, Tang et al., 2016), it appears likely that they have a common functional role in the fine-tuning of synaptic transmission.

### Partial overlap of GlyR and GABA_A_R SSDs at mixed inhibitory synapses

It was shown that most inhibitory synaptic boutons in cultured spinal cord neurons are apposed to PSDs containing both GlyRs and GABA_A_Rs (Dumoulin et al., 2000). Using two-color dSTORM, we therefore compared how GlyRs and GABA_A_Rs were organized at these mixed inhibitory synapses.

In multi-color dSTORM experiments, the choice of fluorescent dyes is one of the main factors determining the quality of the reconstructed super-resolution images (Dempsey et al., 2011, van de Linde et al., 2011). In addition, it is essential to achieve dense labeling, which is very challenging in the case of synaptic proteins, particularly neurotransmitter receptors (discussed in (Yang & Specht, 2020)). We tested different antibody and dye combinations for the detection of GlyR and GABA_A_R clusters in two-color dSTORM. The best combination was to label GABA_A_Rs with secondary antibodies conjugated with Alexa Fluor 647 (A647) and GlyRs with Cy3B (Figure S2), since the number of detections in dSTORM was closely correlated with the cluster intensity in the epifluorescence images (Figure S2A5-B5). The detection efficiency of this dye combination was also relatively good when the amount of receptor labeling was low (Figure S2C1-D5).

Using this labeling protocol, we compared the spatial organization of GlyRs and GABA_A_Rs at mixed inhibitory synapses in cultured spinal cord neurons. Receptor clusters in rendered dSTORM images were highly variable, even though the shape and size of the overall synaptic structure was similar (Figure 3A-B upper panels). GlyRs and GABA_A_Rs were not homogeneously distributed at the PSDs and displayed limited co-localization. To evaluate the level of overlap between the receptor domains, we calculated the overlapped fraction that is the area containing both GlyRs and GABA_A_Rs, divided by the total cluster area (Figure 3C). The overlapped fraction was generally well below 50%. Furthermore, we segmented synaptic clusters into sub-synaptic domains. GlyR SSDs and GABA_A_R SSDs did not always overlap in the binary images (Figure 3A-B lower panels). As judged by the overlap fraction, more than half of the synapses (55%) did not show any co-localization between GlyR SSDs and GABA_A_R SSDs (Figure 3D). In the remainder of the synapses the SSDs of GlyRs and GABA_A_Rs overlapped partially, but were often dominated by one or the other of the two receptors. As a result, the average overlapped fraction of SSD area was only about 9% at these synapses.

**Figure 3.**
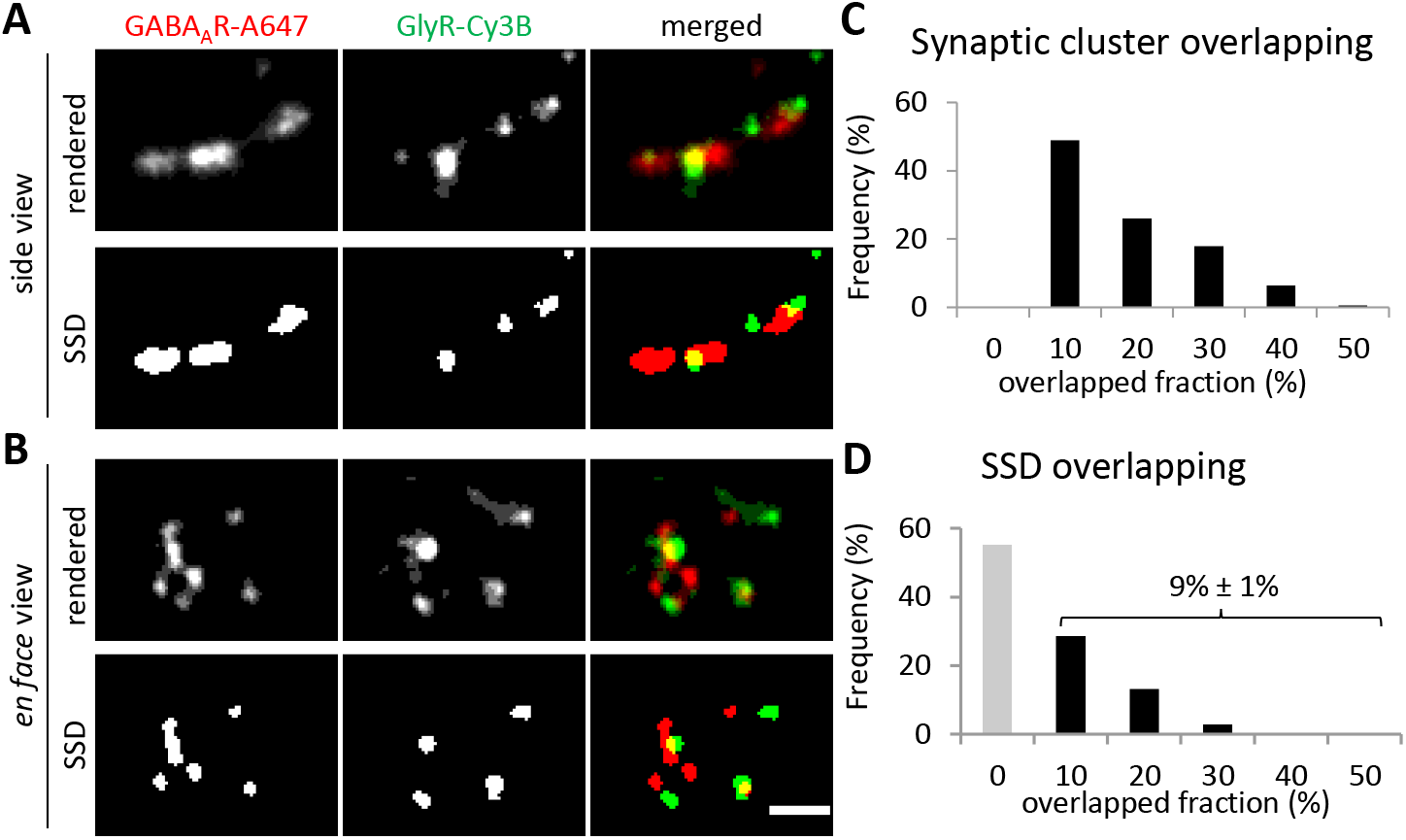
Differential organization of GlyRs and GABA_A_Rs at mixed inhibitory synapses. (A-B) Representative rendered dSTORM images of GlyRs and GABA_A_Rs at synapses in side view (A) or en face view (B). The lower images show the corresponding segmented SSDs. GlyRs were labelled with Alexa 647 and GABA_A_Rs with Cy3B in these experiments. Scale: 200 nm. (C) Quantification of the overlap between GlyRs and GABA_A_Rs at synapses. dSTORM images of GlyRs and GABA_A_Rs were binarized, and the overlapped fraction relative to the total area occupied by the receptors was calculated for each synapse. (D) Overlap between the SSDs of GlyRs and GABA_A_Rs. The fraction of SSD overlap was calculated based on the binarized images for each synapse. Over half of the synapses did not show any overlap between GlyR SSDs and GABA_A_R SSDs (grey bar). The average overlap for the remaining synapses was 9% ±1% (mean ± SEM; n = 395 synapses from three independent experiments, without selection for synapse orientation).

The low level of overlap between GlyR SSDS and GABA_A_R SSDs was unexpected, since both receptors were well aligned with RIM1/2 SSDs (0.79 ± 0.04 and 0.78 ± 0.07, respectively). Consequently, the overlap between GlyR and GABA_A_R SSDs should have been around 62% (0.79 * 0.78 *100%). This apparent discrepancy can be attributed to several factors. Firstly, the synapses in this experiment were not chosen based on their orientation. Instead, this analysis was performed on hundreds of synapses, regardless of whether they were seen in side view or *en face*. Second, the labeling of GlyRs at an extracellular epitope and of GABA_A_Rs at an intracellular epitope may increase the distance between the fluorophore detections in the two channels, thus lowering the apparent overlap. Nevertheless, it appears that GlyRs and GABA_A_Rs form distinct SSDs that are often separated or dominated by one of the two receptor types.

### Differential regulation of GlyRs and GABA_A_Rs in response to altered network activity

Since GlyRs and GABA_A_Rs occupy different sub-synaptic compartments, we hypothesized that they may be controlled through different clustering mechanisms. To investigate whether GlyRs and GABA_A_Rs were differentially regulated by neuronal activity, we adopted a pharmacological approach. Spinal cord neurons were treated either with tetrodotoxin (TTX) or with 4-aminopyridine (4-AP) to reduce or increase the neuronal network activity, respectively. We observed few calcium signals in neurons under control conditions, suggesting a low basal activity in cultured spinal cord neurons.

TTX completely abolished the calcium signals, while 4-AP increased the firing frequency but did not change the amplitude of the calcium transients (Figure S3).

To examine whether GlyRs and GABA_A_Rs were affected by changes in network activity, neurons were treated with TTX and 4-AP for one hour, then fixed and triple labelled with antibodies against GlyRs, GABA_A_Rs and gephyrin. Epifluorescence images in the three channels were taken with conventional fluorescence microscopy (Figure 4A). Synaptic clusters were detected by generating masks based on the gephyrin labeling. The size of the gephyrin puncta (area in pixels) did not differ between the TTX and 4-AP conditions (Figure 4B). The gephyrin masks were then used to measure the intensities of the different associated synaptic receptors. Interestingly, GABA_A_R but not GlyR intensity was significantly decreased after 4-AP treatment, compared to the TTX condition that served as control condition in these experiments (Figure 4C, Figure S4). In order to directly compare how the two receptors reacted at the same PSDs, we calculated the ratio of GABA_A_R/GlyR intensity for each synapse. Lower GABA_A_R/GlyR ratios under 4-AP confirmed that higher network activity decreases synaptic GABA_A_R levels but had no or little effect on GlyRs at the same PSDs (Figure 4D). To further investigate these changes in relation to the sub-synaptic organization of GlyRs and GABA_A_Rs, we then conducted super-resolution imaging in neurons exposed to altered network activity.

**Figure 4.**
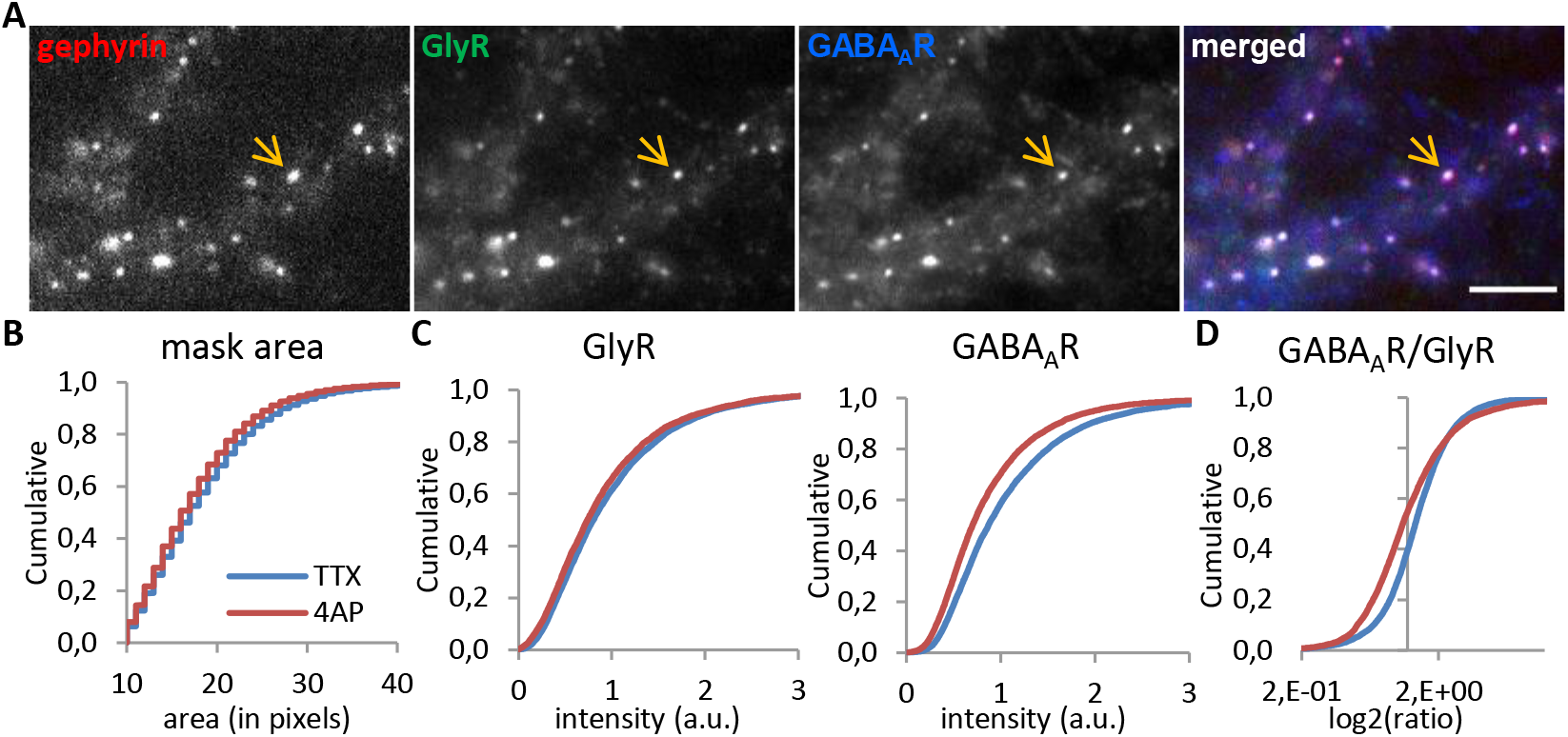
Reduced GABA_A_R levels but not GlyRs at mixed inhibitory synapses by 4-AP treatment. (A) Triple immuno-labeling of gephyrin, GlyRs and GABA_A_Rs at mixed inhibitory synapses (arrow) imaged with conventional fluorescence microscopy. Scale: 5 μm. (B) Quantification of the apparent synaptic area (in pixels) of gephyrin clusters (KS test p < 0.0001). (C) Lower fluorescence intensity of GABA_A_Rs but only a minor reduction of GlyRs at synaptic gephyrin clusters (KS test, p < 0.0001, both panels). (D) Reduced GABA_A_R/GlyR intensity ratio at mixed inhibitory synapses (KS test, p < 0.0001). Number of synapses: n = 9416 in TTX and n = 6949 in 4-AP conditions from three independent experiments.

### Activity-dependent changes of the sub-synaptic organization of GlyRs and GABA_A_Rs

Two-color dSTORM imaging of GlyRs and GABA_A_Rs was performed after one hour of TTX or 4-AP bath application. In order to compare the number of detections of GlyRs and GABA_A_Rs within the same synapse, the rendered dSTORM images of GlyR and GABA_A_R clusters were combined to produce total synaptic clusters that we refer to as *integrated receptor clusters* (Figure 5A). These clusters were binarized to produce a mask of the overall synapse, within which the number GlyR and GABA_A_R detections were determined (Figure 5B). After 4-AP treatment, the number of GABA_A_R detections was significantly decreased, while that of GlyRs did not change (Figure 5C-D). In agreement with the data shown in Figure 4, the ratios of GABA_A_R/GlyR detections per synapse were reduced, confirming that this phenomenon occurred within individual PSDs (Figure 5E). We also saw a reduction in the area of the integrated receptor clusters following 4-AP treatment (Figure S5A). Together, these results established that the copy numbers of GABA_A_Rs and GlyRs at synapses are differentially regulated by network activity.

**Figure 5.**
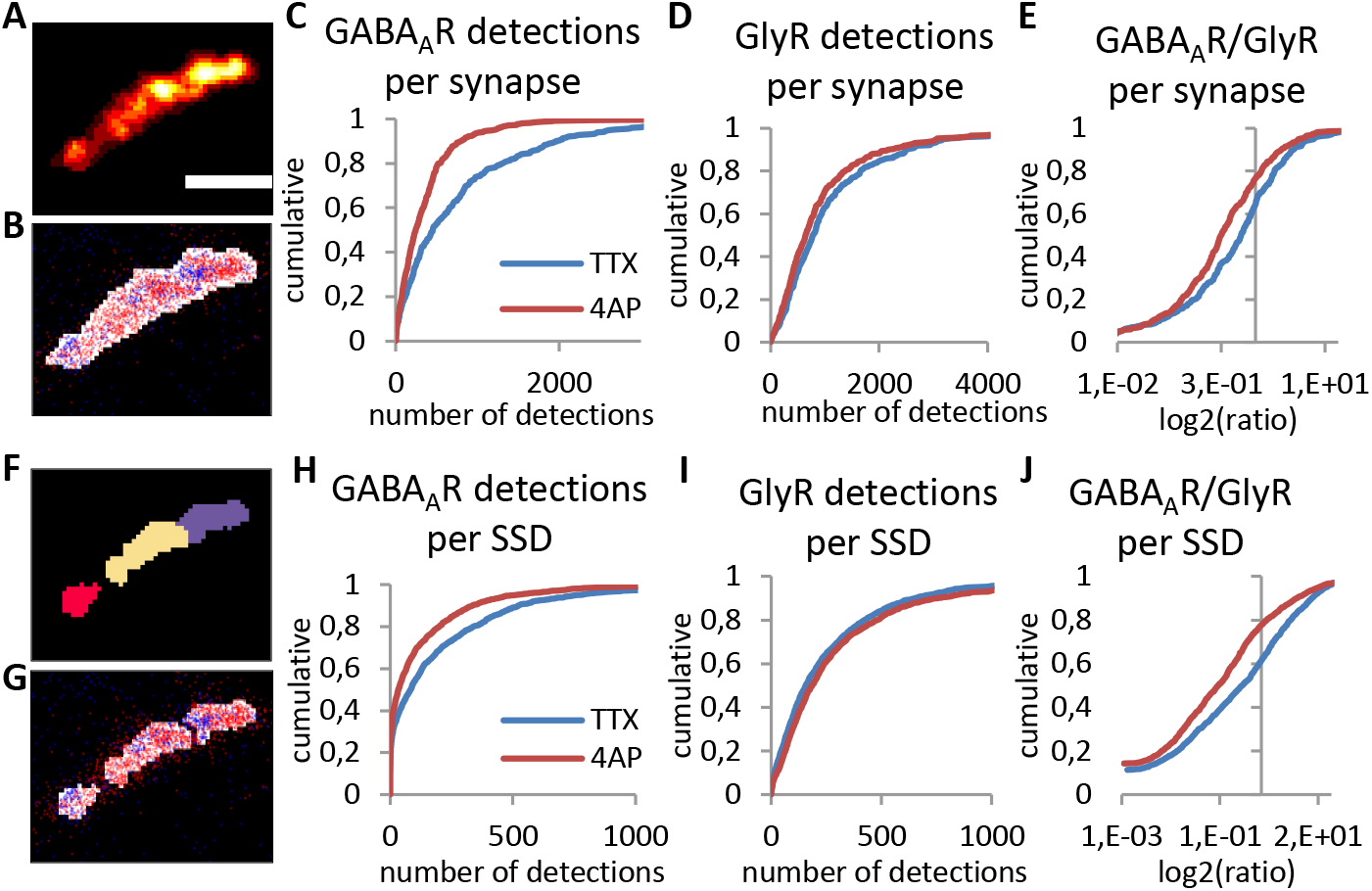
Differential sub-synaptic re-organization of GABA_A_Rs and GlyRs by 4-AP treatment. (A-E) Characterization of the nanoscale organization of GABA_A_Rs and GlyRs at mixed inhibitory synapses. (A) Synaptic receptor cluster rendered from detections of both GABA_A_Rs (labeled with Alexa 647) and GlyRs (labeled with Cy3B). (B) Detections of GABA_A_Rs (red dots) and GlyRs (blue dots) overlaid with binary synaptic mask (white). (C-E) Detections of GABA_A_Rs, but not GlyRs were strongly decreased at mixed synapses after 4-AP treatment (KS test, p < 0.0001 in C and E, p < 0.001 in D). (F-J) Sub-synaptic characterization of GABA_A_R and GlyR domains. (F) Watershed segmentation of subsynaptic domains (SSDs). Each SSD is shown in different colors. (G) Detections of GABA_A_Rs (red dots) and GlyRs (blue dots) overlaid with SSDs (white). (H-J) Detections of GABA_A_Rs but not GlyRs at the same SSDs were reduced after 4-AP treatment (KS, p < 0.0001 in H and J, p < 0.01 in I). Number of synapses: n = 439 in TTX, n = 531 in 4-AP condition, from three independent experiments. Scale: 200 nm.

Next, we segmented the integrated receptor clusters into SSDs, referred to as *receptor SSDs*, and counted the number of detections of GlyRs and GABA_A_Rs within each of them (Figure 5F-G). This allowed us to compare the relative changes of GlyRs and GABA_A_Rs within the same sub-synaptic domains. The size of the receptor SSDs was not affected by the treatments (Figure S5B). However, the number of GABA_A_R detections was decreased by 4-AP, whereas the number of GlyR detections did not change (Figure 5H-I). The reduction of GABA_A_R relative to GlyR copy numbers occurred within the same SSDs, as shown by the ratio of detections per SSD (Figure 5J). These results show that the activity-dependent re-distribution of GABA_A_Rs versus GlyRs occurs also at the sub-synaptic level.

The spatial organization at mixed inhibitory synapses was further assessed by quantifying the number of GlyRs and GABA_A_Rs SSDs separately (Figure 6). The number of integrated receptor SSDs per synapse was reduced after 4-AP treatment (Figure 6A), but not their size (Figure S5B). This reduction could be attributed to a large extent to GABA_A_Rs rather than GlyRs, since the number of GABA_A_R SSDs was strongly reduced (49% of the TTX condition, p < 0.0001, Mann-Whitney test), compared to that of GlyR SSDs (88% of TTX condition, p < 0.01) (Figure 6B-C). This further indicates that the spatial organization of inhibitory receptor domains of GABARs and GlyRs is differentially regulated at mixed inhibitory synapses.

**Figure 6.**
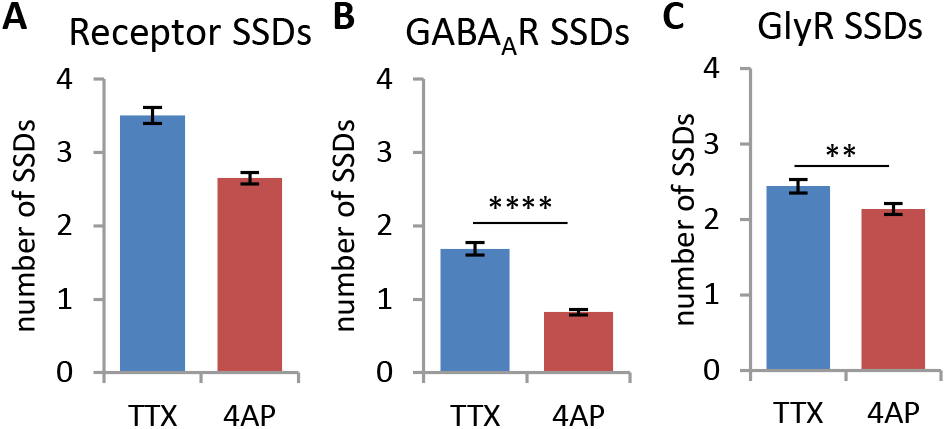
Reduced number of GABA_A_R SSDs after 4-AP treatment. (A) The number of receptor SSDs, segmented from the combined receptor clusters of GABA_A_Rs and GlyRs, was decreased after 4-AP treatment (MW test, p < 0.0001). (B) This was mostly due to the loss of GABA_A_R SSDs that were segmented from the rendered images of GABA_A_R detections only (MW, p < 0.0001). (C) The number of GlyR SSDs (based on GlyR detections) were only marginally reduced (MW, p < 0.01). Number of synapses: n = 452 in TTX, n = 583 in 4-AP condition, from three independent experiments. These analyses are based on the same data set as in Figure 5.

### Sub-synaptic regulation of gephyrin phosphorylation

In order to explore a possible mechanism by which synaptic activity could differentially regulate GlyRs and GABA_A_Rs at mixed inhibitory synapses, we studied the phosphorylation of gephyrin at the regulatory site S270 (reviewed in (Alvarez, 2017, Specht, 2020)). We took advantage of the fact that the monoclonal antibody mAb7a specifically recognizes phosphorylated gephyrin (Kuhse et al., 2012). Hence we examined how this post-translational modification responded to neuronal activity changes.

We first compared the amount of total gephyrin (probed by the polyclonal antibody rbGPHN) and of S270 phosphorylated gephyrin (probed by mAb7a) using conventional fluorescence microscopy (Figure 7A-C). The colocalization of rbGPHN and mAb7a puncta did not differ between the TTX and 4-AP conditions (Figure S6A). Using a binary mask of rbGPHN clusters, we measured the fluorescence intensity of both rbGPHN and mAb7a signals at the same puncta. The fluorescence intensity of mAb7a but not rbGPHN was significantly decreased by 4-AP treatment (Figure 7B, also see Figure S4A). This was also true at individual synapses, as indicated by the fluorescence intensity ratio (Figure 7C). The unchanged total number of gephyrin molecules at synapses suggests that the gephyrin scaffold persists during changes in network activity, while the S270 phosphorylation is dynamically regulated. We therefore explored with dSTORM the sub-synaptic distribution of S270 phosphorylated gephyrin after 4-AP treatment. The number of mAb7a detections per synapse was decreased by 4-AP (Figure 7D), as expected from the reduction of mAb7a labeling seen with conventional fluorescence microscopy (Figure 7B). At the sub-synaptic level, neither the number of detections of pS270 gephyrin (mAb7a) per SSD nor the number of SSDs were altered following 4-AP treatment (Figure 7E-F). Moreover, the size of pS270 gephyrin SSDs did not change (Figure S6B). It thus appears that pS270 gephyrin is concentrated within regions of the synaptic scaffold that are not affected by 4-AP treatment, suggesting that S270 de-phosphorylation occurs predominantly at gephyrin molecules located outside of these SSDs.

**Figure 7.**
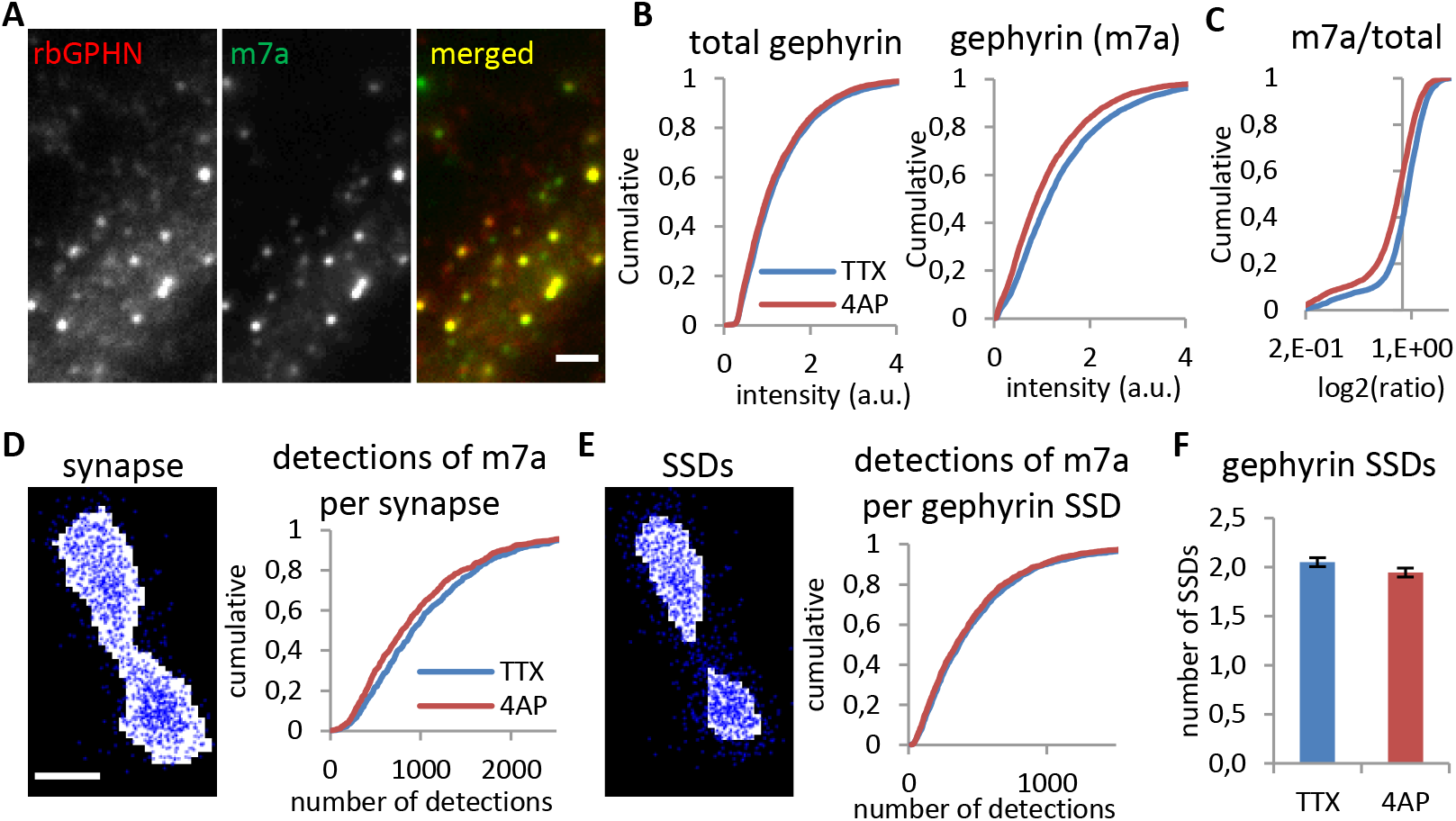
Reduction of gephyrin phosphorylation at synapses but not within SSDs by 4-AP. (A-C) Reduced immuno-reactivity of S270 phosphorylated gephyrin but not total gephyrin levels after 4-AP treatment, revealed by conventional fluorescence microscopy (KS test, p < 0.001 in B1 and p < 0.0001 in B2, p < 0.0001 in C). Total gephyrin was probed with polyclonal rabbit primary antibody (rbGPHN), and pS270 phosphorylated gephyrin with monoclonal mouse primary antibody (m7a). Number of synapses: n = 4040 in TTX and n = 3818 in 4-AP conditions from two independent experiments. Scale: 2 μm. (D-F) Reduced numbers of pS270 gephyrin (m7a) detections were recorded by dSTORM for the entire synaptic area (D, KS test, p < 0.01). The number of detections of pS270 gephyrin per SSD (E, KS, p = 0.18) and the number of SSDs (F, MW, p = 0.11) were not changed by 4-AP treatment. Gephyrin was probed with mAb7a antibody and Alexa 647 dye. Number of synapses: n = 810 in TTX and n = 727 in 4-AP conditions from three independent experiments. Scale bar: 100 nm.

## Discussion

The heterogeneity of synaptic protein distribution has been observed in many instances using superresolution imaging with different techniques such as SMLM, STED and SIM (Crosby et al., 2019, MacGillavry et al., 2013, Nair et al., 2013, Pennacchietti et al., 2017, Specht et al., 2013, Wegner et al., 2018). The sub-synaptic domain (SSD) has been defined as a sub-compartment within the synaptic complex in which the density of a given protein is higher than in the surrounding area (Yang & Specht, 2019). Here, we have shown that several components of mixed inhibitory synapses in spinal cord neurons form SSDs, including RIM1/2, gephyrin, GlyRs and GABA_A_Rs.

To exclude that an inherent stochasticity of dSTORM imaging could be behind the observation of SSDs, we have considered in our experiments the number of detections per secondary antibody (50.4 ± 3.3 for IgG-Alexa 647, and 47.6 ± 3.3 for IgG-Cy3B, mean ± SEM, see (Yang & Specht, 2020)) and extrapolated this value to estimate the number of sampled synaptic receptors. Accordingly, the number of detected GABA_A_Rs (IgG-Alexa 647) per synapse in the control condition was estimated to be roughly 20. The number of detected GlyRs (IgG-Cy3B) per synapse was ~29. The average number of receptors per SSD was 5 for GABA_A_Rs and 8 for GlyRs in the control. Increased network activity by 4-AP treatment reduced the number of detected GABA_A_Rs to 8 per synapse and to 3 per SSD, without affecting the number of GlyRs (23 per synapse, 8 per SSD). Although these values are lower than the absolute copy numbers at synapses (Liu et al., 2020, Nusser et al., 1997, Patrizio et al., 2017), the efficiency of the immuno-labeling nonetheless allows for an accurate structural description.

We found that pre-synaptic RIM1/2 SSDs and post-synaptic gephyrin SSDs are aligned in trans-synaptic nanocolumns at mixed inhibitory synapses. This arrangement is similar to that observed at excitatory synapses, consisting of RIM1/2 and the post-synaptic scaffold protein PSD-95 (Tang et al., 2016). Moreover, GlyR SSDs and GABA_A_R SSDs were also aligned with the SSDs of RIM1/2. This suggests that GlyRs and GABA_A_Rs are integrated into inhibitory the trans-synaptic nanocolumns. Modeling predicts that the positioning of receptors in front of vesicle release sites can increase the transmission efficacy at excitatory synapses (Haas et al., 2018, MacGillavry et al., 2013). Similarly, the alignment of GlyR and GABA_A_R SSDs with RIM1/2 may ensure an effective inhibitory neurotransmission, and provide a means of regulation by slightly shifting receptors from their positions.

At mixed inhibitory synapses, the overlap between GlyR SSDs and GABA_A_R SSDs was surprisingly low, suggesting that the receptor domains are distinct, overlapping only at a subset of synapses in spinal cord neurons. This pattern is reminiscent of the differential distribution of AMPARs and GluN2A and GluN2B-containing NMDARs at excitatory synapses, where the nanoscale organization of the receptors may regulate access to released glutamate (Kellermayer et al., 2018, MacGillavry et al., 2013). In the case of inhibitory synapses, the distribution of GlyRs and GABA_A_Rs SSDs could have a similar impact on their activation, depending on which SSDs are closest to the vesicle release sites, as suggested by the detection of purely glycinergic and GABAergic mIPSCs in young spinal cord neurons (Keller et al., 2001). However, the situation could be more complicated at mixed inhibitory synapses, since GlyRs and GABA_A_Rs are activated by different neurotransmitters, both of which are released from the same pre-synaptic vesicles (Aubrey & Supplisson, 2018, Jonas et al., 1998). The relative vesicle load and binding affinity of the neurotransmitters therefore add to the complexity related to the nanoscale organization that controls the distance of GlyRs and GABA_A_Rs to the release site.

It is generally accepted that gephyrin molecules are organized into a relatively stable lattice at inhibitory PSDs (Alvarez, 2017, Sola et al., 2004). We found that the gephyrin SSDs contain high levels of phosphorylated gephyrin molecules (pS270; Figures 2 and 7). SSDs of pS270 gephyrin have also been detected in previous studies (Crosby et al., 2019, Niwa et al., 2019, Pennacchietti et al., 2017). This raises the concept that the ratio between total and phosphorylated pS270 gephyrin is a key regulatory switch at inhibitory synapses. Given that gephyrin phosphorylation was reduced in response to 4-AP treatment without altering total gephyrin levels, it is possible that pS270 gephyrin is concentrated within sub-domains of the gephyrin scaffold at mixed inhibitory synapses. This is confirmed by the observation that the sub-synaptic pS270 distribution does not change with 4-AP (Figure 7). In other words, the loss of pS270 signal appears to occur in regions of the PSD that do not form part of the SSDs. The relative stability of gephyrin phosphorylation inside and out of SSDs may thus provide a potential mechanism for the differential regulation of GlyR and GABA_A_R clustering at mixed inhibitory synapses.

When network activity levels were altered, the nanoscale organization of GlyRs and GABA_A_Rs was differentially adjusted (Figures 5 and 6). While GABA_A_R clustering was reduced in response to higher activity after 4-AP treatment, GlyRs remained largely unchanged. GlyR levels may be sustained by the gephyrin scaffold independently of changes of its phosphorylation status due to the high binding affinity of the GlyRβ subunit for gephyrin. In contrast, de-phosphorylation of gephyrin outside of SSDs could be responsible for the homeostatic reduction of GABA_A_Rs under 4-AP. This is in agreement with the reduced GABA_A_R clustering downstream of gephyrin de-phosphorylation induced by cAMP signaling (Niwa et al., 2019). Taken together, our results demonstrate that GlyRs and GABA_A_Rs are differentially regulated by gephyrin through different signaling pathways. The fact that GlyRs and GABA_A_Rs occupy distinct sub-synaptic domains and are regulated by different signaling pathways provides a new angle for the understanding of co-transmission at mixed inhibitory synapses in spinal cord neurons.

## Acknowledgements

We thank Marianne Renner for providing the pickpointsSR program, Philippe Rostaing and Oliver Gemin for preparing cryosections, Felipe Delestro and Auguste Genovesio for technical support in data analysis. Funding was provided by the European Research Council (ERC, Plastinhib), Agence Nationale de la Recherche (ANR, Synaptune and Syntrack), Labex (Memolife) and France-Biolmaging (FBI). XY was supported by the China Scholarship Council.

## The authors declare that they have no conflict of interest

## Methods

### Primary spinal cord neuron culture

All procedures using animals follow the regulations of the French Ministry of Agriculture and the Direction départementale des services vétérinaires de Paris (Ecole Normale Supérieure, animalerie des rongeurs, license B 75-05-20). Primary spinal cord neurons were prepared from embryonic Sprague Dawley rats on embryonic day E14 as described (Specht et al., 2013), with some modifications. Dissociated neurons were plated on 18 mm glass coverslips (thickness 0.16 mm, No. 1.5, VWR #6310153) that were pre-washed with ethanol and coated with 70 μg/ml poly-DL-ornithine. Cells were seeded at a density of 6 × 10^4^ cells/cm^2^ in Neurobasal medium (ThermoFisher), supplemented with B-27, 2 mM glutamine, and antibiotics (5 U/ml penicillin and 5 μg/ml streptomycin). Neurons were cultured at 37°C with 5% CO_2_, and the medium was replenished twice a week by replacing half of the volume with BrainPhys neuronal medium (Stemcell Technologies) supplemented with SM1 and antibiotics. Cells were used for experiments between DIV14 and DIV21.

### Calcium imaging

Spinal cord neurons were loaded with 0.5 μM Fluo-4 AM (F14201, Life Technologies) diluted in culture medium and incubated for 10 min. Cells were then imaged within pre-warmed imaging medium, containing 130 mM NaCl, 5 mM KCl, 2 mM CaCl_2_, 1 mM MgCl_2_, 30 mM glucose, and 10 mM HEPES, pH 7.4. Time lapse recordings were acquired for 3 min at 10 Hz before and after application of TTX (1 μM) or 4-AP (50 μM), respectively. Background corrected calcium signals were measured in cell bodies using Fiji (Schindelin et al., 2012). The frequency and peak amplitude was detected using a custom written program (courtesy from Anastasia Ludwig) in Matlab (MathWorks).

### Pharmacological treatment and immunocytochemistry

Cultured neurons were treated for one hour with tetrodotoxin citrate (TTX, Cat. No. 1069, Tocris) or 4-aminopyridine (4-AP, Cat. No. 0940, Tocris), diluted in culture medium at a final concentration of 1 μM and 50 μM, respectively. The cultures were then fixed with 100% methanol at −20°C for 10 minutes. Unspecific binding sites were blocked with 3% BSA in PBS (blocking buffer) for at least 30 minutes. Cells were sequentially incubated for 1 hour at room temperature with primary antibodies and with secondary antibodies diluted in blocking solution. Each step was followed by several washes with PBS.

The following primary antibodies were used: mouse monoclonal mAb7a (m7a, Synaptic Systems, #147011), rat monoclonal mAb7a (rb7a, Synaptic Systems, #147208), and rabbit polyclonal (rbGPHN, Synaptic Systems, #147002) antibodies against gephyrin; mouse monoclonal antibody against GABA_A_R β3 (UC Davis/NIH NeuroMab, #75-149, RRID: AB_2IO9585), rabbit polyclonal antibody against RIM1/2 (Synaptic Systems, #140203), guinea pig polyclonal antibody against RIM1/2 (Synaptic Systems, #140205), and custom-made rabbit polyclonal antibody against GlyR α1 (Triller lab, #2353). The primary antibodies were used at a dilution of 1:500, following the manufacturer’s instructions. The following commercial secondary antibodies were used: Alexa Fluor 647-conjugated donkey anti-guinea pig IgG (Jackson ImmunoResearch, Cat. No. 706-605-148), donkey anti-rabbit (Jackson, Cat. No. 711-605-152), donkey anti-mouse (Jackson, Cat. No. 715-605-151) and goat antirat IgG (Invitrogen, Cat. No. A21247); Cy3-conjugated goat anti-rabbit (Jackson, Cat. No. 111-165-144) and goat anti-mouse IgG (Jackson, Cat. No. 115-165-166); Alexa 488-conjugated goat antimouse IgG (Jackson, Cat. No. 115-545-166). Secondary antibodies were diluted at 1:500 or 1:1000, following the manufacturers’ recommendations. Secondary antibodies that had been conjugated with Cy3B in our lab were diluted at 1:50 or 1:100 to compensate for the dilution during the purification process.

### Conjugation of secondary antibodies with Cy3B dye

Unconjugated secondary antibodies, donkey anti-mouse IgG (Jackson ImmunoResearch, #715-005-151), donkey anti-rabbit IgG (Jackson ImmunoResearch, #711-005-152), donkey anti-guinea pig IgG (Jackson ImmunoResearch, #706-005-148) were coupled with Cy3B mono-reactive NHS ester (PA63101, GE Healthcare) according to the supplier’s protocol. Antibodies were then purified using size exclusion columns (Illustra NAP-5 columns, #17085302, GE Healthcare). The absorption of IgG at 280 nm and Cy3B dye at 559 nm of the reaction products was measured by spectrophotometry (NanoDrop ND-1000 Spectrophotometer), from which the number of dyes per IgG was calculated. The estimated dye/IgG ratio was generally between 3 and 5.

### Cryosection preparation and immunohistochemistry

Adult male mice (C57BL/6J, 10 weeks old) were deeply anesthetized with pentobarbital, and intracardially perfused with 4% PFA and 0.1% glutaraldehyde in PBS. The cervical to thoracic spinal cord segments were dissected and post-fixed in 4% PFA at 4°C overnight. The segments were then cut into 1 mm^3^ cubes and incubated in 2.3 M sucrose at 4°C for 2-3 days. The tissue was then cut into sections of 2 μm thickness with an ultra-microtome (Leica EM UC6). Sections were placed on glass coverslips for immunohistochemistry.

Sucrose impregnated cryosections were first subjected to a heat-induced antigen retrieval (HIAR) protocol (Rousseau et al., 2012). Briefly, sections were immersed in a citrate-based antigen retrieval buffer and placed in a de-cloaking chamber (Biocare Medical) for 20 min at 110°C. They were then treated with 15% methanol and 0.3% H_2_O_2_ in PBS for 15 min, followed by 1% sodium borohydride in PBS for 45 min. Sections were thoroughly rinsed with PBS after each step. Unspecific binding sites were blocked with 3% BSA in PBS at room temperature for at least two hours. Primary antibodies, including rabbit polyclonal antibody against RIM1/2 and mouse monoclonal mAb7a, were diluted in blocking buffer and applied overnight at 4°C. Sections were then incubated with Alexa 647-conjugated donkey anti-rabbit IgG (Jackson ImmunoResearch, Cat. No. 711-605-152, dilution 1:500) and lab-made Cy3B-conjugated donkey anti-mouse secondary antibodies (dilution 1:100) for two hours at room temperature.

### Conventional fluorescence microscopy and image analysis

After immuno-labeling, coverslips were mounted in an open chamber in PBS. Images were acquired on an inverted Nikon Eclipse Ti microscope, equipped with a 100× oil immersion objective (HP APO TIRF 100× oil, NA 1.49, Nikon). A mercury lamp (Intensilight C-HGFIE, Nikon) was used for illumination, with specific band pass filters for the far-red (excitation FF01-650/13, emission FF02-684/24, Semrock), red (ex. FF01-560/25, em. FF01-607/36) and green channels (ex. FF02-485/20, em. FF01-525/30). Ten image frames of 100 ms or 200 ms exposure (depending on the channel) were recorded for each field of view. For a given set of experiments, all imaging parameters were kept constant between all experimental conditions. The stacks of images were combined by average projection of the ten frames in Fiji. Binary masks were then produced from the gephyrin immunolabelling using the wavelet function in Icy (de Chaumont et al., 2012), to measure the intensities of all the different labeled proteins within the synaptic cluster. The integrated intensity within the binary masks was determined using a lab-made program (Hennekinne et al., 2013) written in Matlab.

### Sequential two-color dSTORM imaging

The dSTORM setup is built on the same Nikon Ti microscope described above. It includes several continuous laser lines with emission wavelengths at 640 nm, 561 nm and 405 nm (Coherent) with nominal maximum power of 1W, 1W, and 120 mW, respectively. The setup is equipped with a total internal reflection fluorescence (TIRF) arm to set the angle of the illumination laser, an acousto-optic tunable filter (AOTF) to control the intensity of the lasers, a perfect focusing system (PFS) to maintain the focal plane during imaging, and an electron multiplying charge-coupled device (EMCCD) camera (Andor iXon Ultra) for image acquisition. All elements of the microscope are controlled by NIS-Elements software (Nikon). The dSTORM recordings were taken at a magnification of 100x, resulting in image pixel size of 160 nm.

On the day of imaging, immuno-labeled neurons on glass coverslips were incubated with 100 nm beads (Tetraspeck, ThermoFisher, Cat. No. T7279), and mounted on glass slides with a cavity (diameter 15-18 mm, depth 0.6-0.8 mm) containing freshly prepared Gloxy imaging buffer (0.5 mg/ml glucose oxidase, 40 μg/ml catalase, 0.5 M D-glucose, 50 mM β-mercaptoethylamine (MEA) in PBS, pH 7.4, degassed with nitrogen). The coverslips were sealed with silicone rubber (Picodent twinsil speed 22) and the slide was mounted on the microscope. Prior to dSTORM recordings, conventional epifluorescence images of each field of view were taken with the mercury lamp (Figure S2A1-D1). These epifluorescence images were used as reference during the dSTORM data processing. Sequential two-color dSTORM imaging was then carried out first in the far-red channel (Alexa 647), followed by the red channel (Cy3B). No UV light was applied during the recording of Alexa 647, however, the blinking of Cy3B was supported by low-intensity 405 nm laser illumination. Highly inclined illumination was used to reduce the background fluorescence. Recordings of 20000 or 30000 frames with an exposure time of 50 ms per frame were acquired. The frame numbers were kept constant for each set of experiments. For single-color dSTORM imaging, only the signals of Alexa 647 in the far-red channel were collected.

### dSTORM data processing

Processing steps of the imaging data included single particle detection, lateral drift correction, two-channel registration and image reconstruction (Yang & Specht, 2020). Single particle detection was realized by applying Gaussian fitting to the PSF of each fluorophore signal, using an adapted version of the multiple-target tracing (MTT) program (Serge et al., 2008) in Matlab. We obtained a localization precision of 6 ± 3 nm for Alexa 647 signals, and 7 ± 2 nm for Cy3B, by the approximation of Δ/√N, where Δ is the full width at half-maximum (FWHM) of the point spread function (PSF) and N is the number of collected photons. Lateral drift was corrected in the Matlab program PALMvis (Lelek et al., 2012) using beads as fiducial markers. At this stage, we obtained the coordinates-based localization data in both channels (Figure S2A2-D2). The drift-corrected coordinates were then rendered into super-resolution dSTORM images (pixel size 10 nm) by representing each localization as a Gaussian function with a standard deviation σ = 15 nm (Figure S2A3-D3). Image rendering was done either with PALMvis to obtain a well-defined structural representation or with LAMA software (Malkusch & Heilemann, 2016) if a consistent representation of detection numbers was required.

### SSD segmentation and feature extraction

To define the regions of interest (mixed inhibitory synapses), individual synaptic clusters were cropped from the rendered images using binary masks produced from the epifluorescence images. SSDs were segmented from these individual synaptic clusters using the H-watershed plugin developed by Benoit Lombardot (http://imagej.net/Interactive_Watershed) in Fiji. The SSD counts per synaptic cluster and SSD area were extracted using extended particle analyzer (Brocher, 2014) in Fiji.

### Quantification of SSD distances and alignment

Rendered dSTORM images of the two proteins of interest were composed and transformed to RGB format in Fiji. SSD distances and alignment were only measured in synapses seen in cross-section. We manually selected these side views following the following criteria: clusters showing obvious elongated shapes in both channels, or synapses with elongated shapes in only one of the two channels and with clear signals in the other channel. After SSD segmentation, the paired SSDs of two different proteins were identified. In the RGB images, a line was drawn through the intensity peaks of the paired SSDs and the distance between the peaks was taken as SSD distance. When there was more than one pair of SSDs per synapse, the average of all measured distances was taken as SSD distance for that synapse. Paired SSDs of pre-synaptic and post-synaptic proteins were counted manually. For each protein, the degree of alignment was calculated as the number of paired SSDs divided by the total number of SSDs for that protein, values expressed on a scale from 0 to 1 (fraction of aligned SSDs).

### Counting dSTORM detections per synapse and per SSD

To obtain the binary masks of synaptic area, the rendered images from the two dSTORM channels generated with LAMA software were combined and binarized without thresholding. For one-color dSTORM, the rendered images were binarized directly to obtain synaptic masks. Binary SSD images were produced with the H-watershed plugin in Fiji. The detection coordinates were then overlaid with the binary images and detections within the synaptic masks or SSDs were counted using the pickpointsSR program (courtesy from Marianne Renner, IFM, Paris) written in Matlab (Figure S2A4-D4).

### Statistics

Statistical analysis was generally done using the non-parametric Mann-Whitney U-test (MW), Kolmogorov-Smirnov test (KS), and Friedman test followed by a Dunn’s post-hoc multiple comparison test (in Figure S3C). Where the data passed a D’Agostino & Pearson normality test, the analysis was done with a Student’s t-test or one-way ANOVA followed by Tukey’s post-hoc test. Data are represented in the form of cumulative probability distributions, or as bar graphs showing the mean ± SEM (or the median and the quartiles of the distribution in Figure S3) unless otherwise stated.

